# Candida albicans exhibits distinct cytoprotective responses to anti-fungal drugs that facilitate the evolution of drug resistance

**DOI:** 10.1101/2020.01.21.914549

**Authors:** V Bettauer, S Massahi, S Khurdia, ACBP Costa, RP Omran, N Khosravi, S Simpson, M Harb, V Dumeaux, M Whiteway, MT Hallett

**Affiliations:** Department of Biology, Concordia University, Montreal, Quebec, Canada; Centre for Applied Synthetic Biology, Concordia University, Montreal, Quebec, Canada; Department of Computer Science and Software Engineering, Concordia University, Montreal, Quebec, Canada; PERFORM Centre, Concordia University, Montreal, Quebec, Canada

**Author notes:** These authors contributed equally to the manuscript (V Bettauer, S Massahi).

## Abstract

We developed a modified protocol for nanolitre droplet-based single cell sequencing appropriate for fungal settings, and used it to transcriptionally profiled several thousands cells from a prototrophic *Candida albicans* population and several drug exposed colonies (incl. fluconazole, caspofungin and nystatin). Thousands of cells from each colony were profiled both at early and late time points post-treatment in order to infer survival trajectories from initial drug tolerance to drug resistance. We find that prototrophic *C. albicans* populations differentially and stochastically express cytoprotective epigenetic programs. For all drugs, there is evidence that tolerant individuals partition into distinct subpopulations, each with a unique survival strategy involving different regulatory programs. These responses are weakly related to changes in morphology (shift from white to opaque forms, or shift from yeast to filamentous forms). In turn, those subpopulations that successfully reach resistance each have a distinct multivariate epigenetic response that coordinates the expression of efflux pumps, chaperones, transport mechanisms, and cell wall maintenance. Live cell fluorescent imaging was used to validate predictions of which molecular responses most often led to survival after drug exposure. Together our findings provide evidence that *C. albicans* has a robust toolkit of short-term epigenetic cytoprotective responses designed to “buy time” and increase the chance of acquiring long-term resistance.

## Introduction

*Candida albicans* is both a human commensal and opportunistic pathogen. Nosocomial infections due to pathogenic *C. albicans* are the fourth most important in North America and are associated with significant socioeconomic burden^1,2^. Systemic Candida infections of immune-compromised individuals are frequently lethal even when treated optimally^3^.

There are at least five classes of anti-fungal drugs: the polyenes, the azoles, the allylamines, the echinocandins, and the nucleoside analog flucytosine. The first four attack a component of the pathogenic fungi that is distinct from the human host^4^: the polyenes cause membrane leakage through interaction with ergosterol whereas the azoles and allylamines block the synthesis of ergosterol at different steps and the echinocandins attack the biosynthesis of the fungal cell wall. Flucytosine is metabolized into compounds that interfere with fungal DNA replication and RNA production; it is typically used in combination with the polyene amphotericin B.

The interaction between *C. albicans* and antifungal drugs is complex and multiple factors determine how the pathogen will be cleared or persist in the host individual. For example, persistence may be due to defects in the immune system of the host^5,6^ or due to pre-existing or acquired genetic mechanisms of genetic resistance^7^. For example, flucytosine resistance can result from mutations in the cytosine permease leading to a block in drug uptake, or by mutations in cytosine deaminase or uracil phosphoribosyltransferase that prevent the compound from being turned into a toxic analog^8^. Resistance to azoles can result from increased drug efflux^9–11^ or mutations in the drug target *ERG11*^12^. Similarly, terbinafine resistance can arise from mutations in the drug target squalene epoxidase^13^, while echinocandin resistance occurs through mutations in its target, the 1-3 beta glucan synthase^14^. Mutations in other elements can influence resistance more indirectly. For example, mutations in alternative components of the ergosterol pathway can provide resistance to azoles by preventing the blocked step from generating the highly detrimental sterols the wild type cells produce when Erg11 function is compromised^15^. Stress response pathways, particularly involving Hsp90 circuitry, are also linked to resistance to antifungal compounds^16^. Importantly, recent unbiased genomic profiling of 43 clinical isolates identified loss of heterozygosity events and single nucleotide polymorphisms in over 240 genes involved in adherence, filamentation, virulence and other processes, suggesting that genetic acquired resistance may be achieved in many ways including via genomic instability^17^. There is now a substantial literature suggesting that *C. albicans*, like *S. cerevisiae*, generate large scale genomic variation as a means of adaptation^18–23^, and there is some evidence that this could be facilitated by the parasexuality of the fungus^18,20,24–27^.

*C. albicans* is well adapted to its role as an opportunistic pathogen with several distinct cellular morphologies that predominate in different niches. Drug response, resistance and clinical persistence are interwoven with these *C. albicans* morphologies including the white yeast form implicated in bloodstream dissemination, the opaque yeast form found predominantly in skin infections^28^, and the hyphal form associated with tissue invasion^29^. Each morphology has unique underlying metabolic and regulatory characteristics that play cytoprotective roles against anti-fungals^30^. Molecular signatures for each transition from white to opaque or filamentous morphologies are different and vary according to the anti-fungal, strain or environmental conditions^31–36^.

Together, these molecular and cellular changes in *C. albicans* constitute the drug tolerance response, a short time interval post-treatment that is inherently epigenetic in nature, involving “reversible” regulatory programs and an absence of fixed genetic mutations. That is, the source of the cells physiological ability to survive in the presence of the inhibitory compound is not due to stable genomic modifications, but rests upon post-transcriptional programs such as position in the cell cycle, expression of stress genes, or random fluctuation in key cellular defense genes. In general, tolerance is not nearly as well understood as drug resistance^37,38^. Even when grown under standard laboratory conditions, isogenetic (or near isogenic) Candida populations respond differentially to antifungal drugs, with some fraction of the population exhibiting drug tolerance, defined as the ability of individuals to survive and grow at drug concentrations above the minimal inhibitory concentration (MIC)^37,39^. Bet hedging, for example, in response to environmental stress, has been observed in *S. cerevisiae*^40,41^. Tolerance also appears to be connected with reduced drug accumulation, where tolerant cells appear physiologically more capable of preventing drug uptake or better at drug export. Although time scales are almost certainly longer within *in vivo* contexts, the tolerance phase, defined as the “lag time” between drug introduction and true acquired resistance, is relatively short; in some cases only a 48hr window is sufficient time for the fixation of advantageous genetic lesions^18,20,37^.

The *C. albicans* drug tolerance response “buys cells time” in order to generate *de novo* genetic mutations, thus increasing the chance of long-term survival. However, we do not fully understand these epigenetic programs and how they vary between drugs and environmental conditions. Our effort here is structured around the hypothesis that survival of any individual *C. albicans* cell, which does not already harbour a latent advantageous genetic mutation conferring drug resistance, is able to survive because it achieves an advantageous epigenetic configuration at the time of the drug challenge. This epigenetic configure provides a window for the generation of somatic mutational events for long-term resistance (**Figure 1A**). This effort exploits our novel single cell sequencing approach, the first droplet sequencing (DROP-seq) method applicable for fungi. We identify subcommunities that successfully tolerate different anti-fungal agents with distinct modes of actions. We further investigate their dynamics using live cell imaging and characterize the set of cytoprotective molecular pathways and processes that could be targeted to ablate the tolerance phase, thereby minimizing the likelihood of acquiring long-term drug resistance.

**Figure 1.**
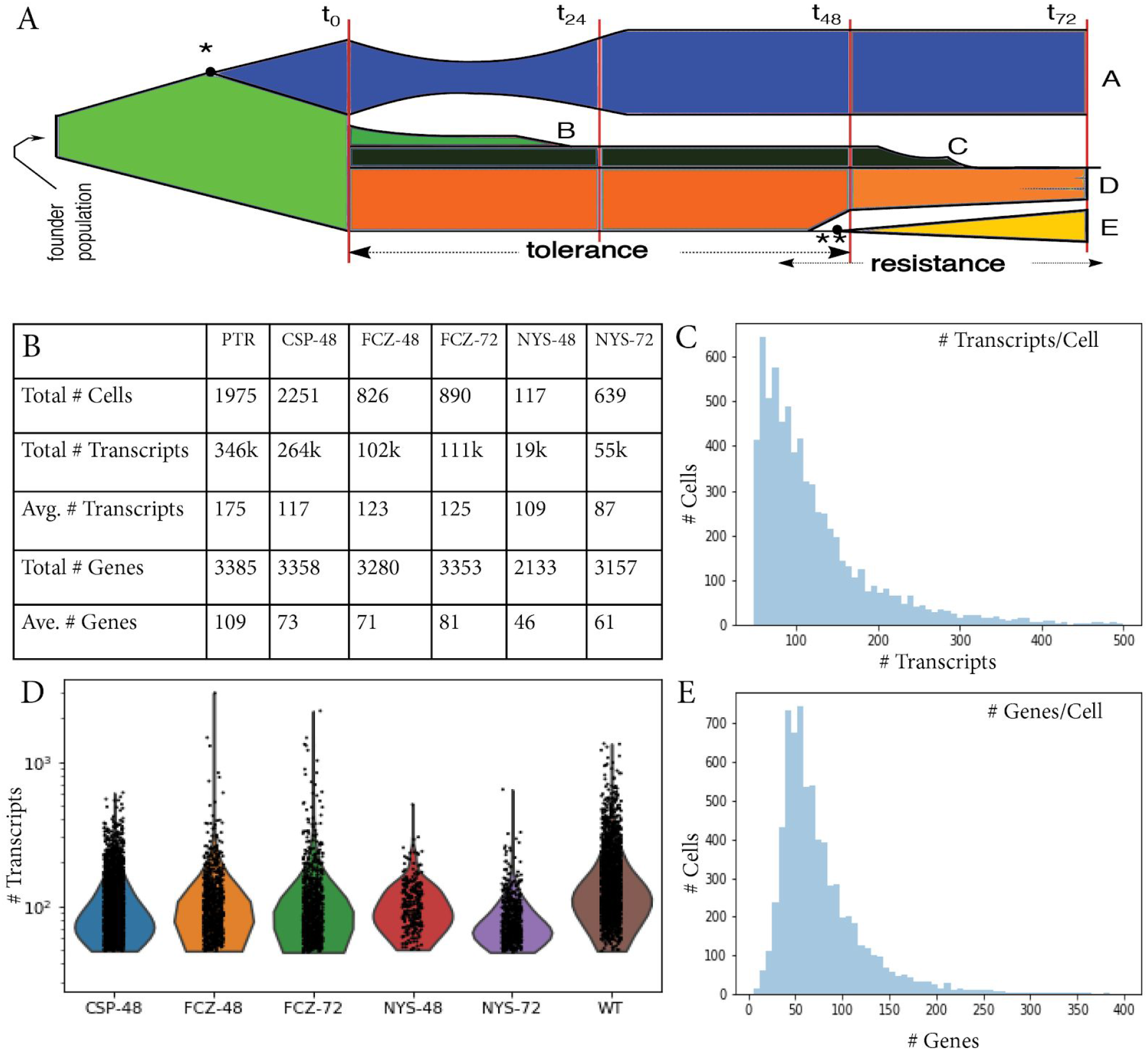
**A.** The potential trajectories of a fungal population challenged with an antifungal agent at time t_0_.The pre-existence or the quick evolution in the founder population of a latent advantageous mutation (**\*** and subsequent blue fraction) before drug exposure confers survival (population A). The remaining populations B-E lack such a preexisting mutation. Cells that do not mount a sufficient epigenetic tolerance defense die off (B). This proposal is focused on populations C,D,E that are each able to enlist an appropriate tolerance response. In some subpopulations (E), genetic events ultimately occur that confer long-term resistance (**\*\*, yellow**), whereas other subpopulations (C) may die off. Some subpopulations (D) may survive the drug through epigenetic modulation alone. **B.** Survey of results from fungal DROP-seq across different populations. **C, E.** Histograms describing the number of cells with observed levels of transcripts and genes respectively. **D.** Violin plots describing the distribution of transcript number.

## Results

### A droplet-based single cell sequencing approach for fungi (fungal DROP-seq)

The *C. albicans* setting required an optimized protocol for cell preparation with specific techniques to remove the cell wall and induce stable spheroplasts, special agents to fix the transcriptome, and filtering steps to separate very large cells with hyphae morphologies. SC5314 populations were grown in YPD media alone (prototrophic, PTR) or in the presence of an antifungal for either 48 or 72 hrs (**Methods 1, 2**). We chose a concentration of 0.01 mg/ml for both fluconazole (FCZ) and nystatin (NYS), representing a moderate dosage relative to their MIC_50_s^8,10,48–50^. For caspofungin (CSP), a compound that interrupts cell wall biosynthesis^51–53^, a concentration of 1 nanogram/ml. This is well below its MIC_50_ levels and chosen in order to increase the number of survivors of this compound (**Methods 3**). All cultures at all time points yielded a sufficient population of survivors for downstream DROP-seq profiling, although 72 hr Caspofungin (CSP-72) was excluded from the present study for logistical reasons. The cells were processed with our fungal DROP-seq device, and the captured material sequenced with the Illumina NEXT-seq following a standard protocol^43,44^ with 200M read/sample (**Methods 4-6**). The raw sequencing data was subjected to our bioinformatics pipeline for read processing, data normalization and imputation (**Methods 7**). **Figure 1B, C** and **Supplemental Figure 1** provide summaries of the observed data post-sequencing, and the effects of normalization and imputation respectively. Results are comparable to previous studies^43^ corrected for the size of the Candida transcriptome.

### Different anti-fungals exhibit distinct transcriptional responses

We asked whether *C. albicans* populations exhibit distinct responses to different anti-fungal drugs. **Figure 2** provides an unsupervised clustering of the single cell expression profiles (UMAP, **Methods 8**) labelled with population of origin (panel **A**) and clusters using VISION (panel **B**). Alternative dimensionality reduction techniques and visualizations^65,66^ produced qualitatively similar cell clusters (data not shown). Although PTR cells are likely near isogenic, they exhibit considerable transcriptional variation, bridging from some late FCZ-72 survivors (cluster 14) to late NYS-72 survivors (cluster 1). FCZ-72 and NYS-72 survivors have distinct gene expression patterns, each separating into two non-overlapping clusters. One subpopulation of CSP-48 and a subpopulation FCZ-48 are highly intermixed in clusters 9, 16, 18 20 (panel **B**). Far from this location, a second pair of subpopulations of CSP-48 and FCZ-48 form cluster 6. In fact, cluster 6 contains cells from almost all drugs and timepoints.

**Figure 2.**
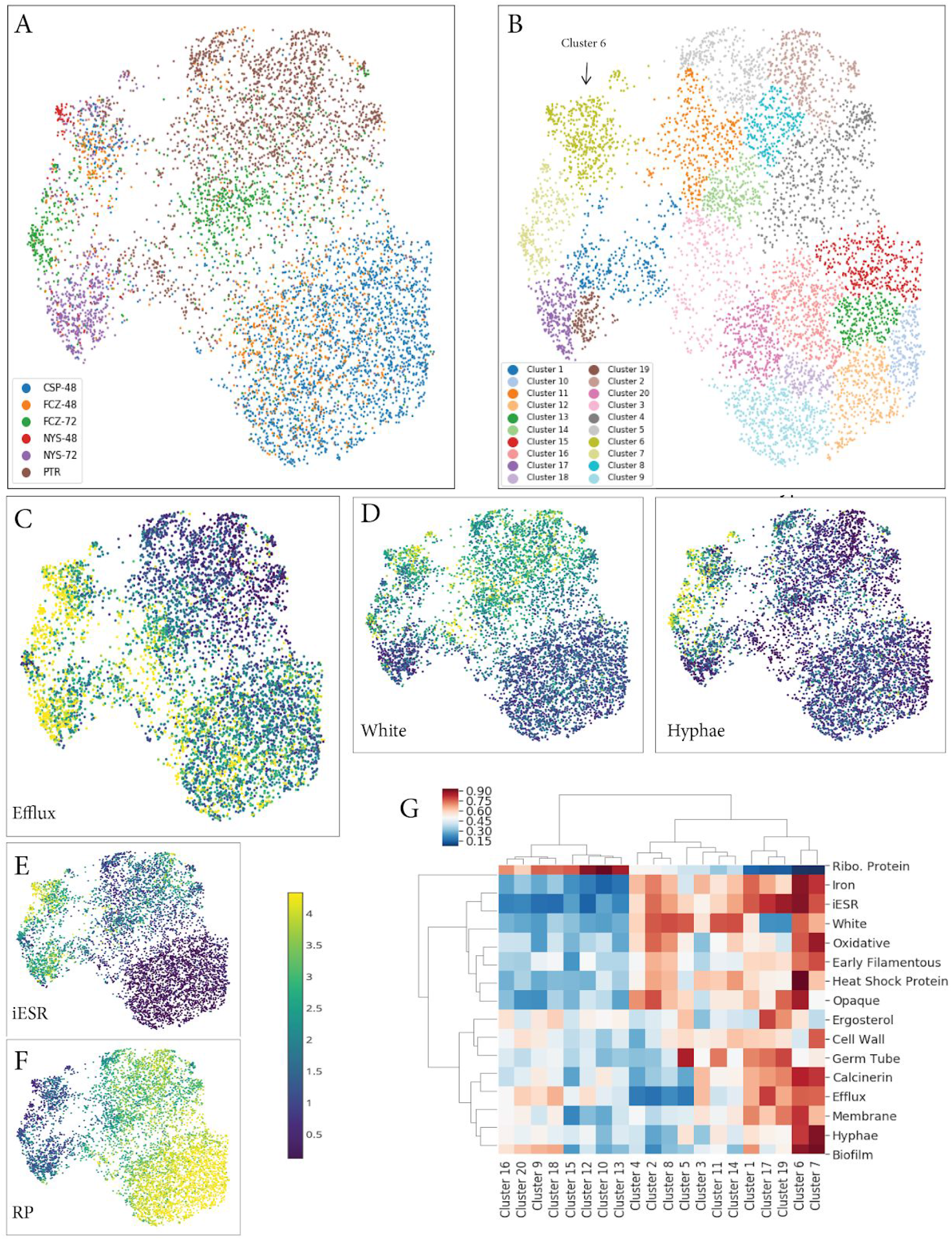
**A.** UMAP based visualization of the relationship between all *C. albicans* populations labelled by population of origin. **B.** UMAP embedding from A but here color reflects unsupervised Louvain found that many have well established roles in different stress responses including HSP21, HGT6 and CAS5 (core stress response), GAC1, XYL2 and ADH2 in acid stress, SOD3, YCF1 and OXR1 in clustering using VISION. **C.** Pattern of expression of the efflux pump gene signature mapped onto the UMAP embedding of A. **D.** Pattern of expression of white and hyphae gene signatures. **E, F.** Pattern of expression for the iESR and Ribosomal Protein (RP) signatures. **G.** Summary of expression of our collection of all signatures across the (unsupervised) Louvain clusters across all cell populations.

We next investigated whether specific processes and pathways were differential between the subpopulations highlighted in panels **A** and **B**. Towards this end, we collected from the literature genes or transcriptional signatures for relevant biological processes (**Methods 9**), and summarized the joint expression pattern of this set of genes for each cell using VISION (**Methods 10**). Drug resistance genes including those coding for efflux pumps were generally expressed lowest in PTR cells **(Figure 2C)** as perhaps expected. Moreover, PTR cells had the most evidence of expressing the white yeast morphology and the least evidence of hyphae transition signatures, compared to drug treated populations (**Figure 2D**). We note however that almost every PTR cell was in the white yeast C. morphology (data not shown).

Many of the genes differentially expressed between PTR and drug treated populations were related to cellular morphology including PFY1 (1.5 logFC), WH11 (1.77), PST1 (2.0). Also supporting the quality of our data, we observed that the inducible Environmental Stress Response (iESR)^40,41^, a signature that should be upregulated in stressed cells, varied significantly across our panel and was inversely correlated with the expression of ribosomal proteins, which tend to be expressed higher in stable healthy cells^40^ (**Figure 2E, F**). We were able to verify that the clusters from Panel **B** are not primarily driven by cell cycle state (**Supplemental Figure 1**). Overall, we observe a varied response for many additional processes (**Figure 2G**). Together this suggests that *C. albicans* mounts distinct molecular responses to different classes of anti-fungal drugs.

### Untreated colonies exhibit significant heterogeneity

We also observed from **Figure 2** that there is significant transcriptional heterogeneity across PTR cells involving at least seven VISION clusters (**Panel B**). Of particular note, mustard cluster 6 is diverse, containing PTR cells and cells from all drug/time points. Dark blue cluster 1 is intermixed with late NYS-72 survivors. Some remaining PTR cells border late FCZ-72 survivors.

In order to better characterize molecular differences here, we restricted attention to only PTR cells, applied Louvain clustering and identified 15 distinct transcriptional regions (**Figure 3A**). These three regions (cluster 8 versus clusters 1, 12 versus clusters 2, 4 and 10) express the iESR. The iESR is in turn broadly anti-correlated with the ribosomal proteins (RP) signature, a marker of healthy cells (**Figure 3B**). Our goal was then to characterize these three regions across our panel of signatures (**Figure 3E**). Cluster 8 strongly expresses most signatures except ergosterol and cell maintenance, suggesting the cells are very active although grown in a prototrophic environment with sufficient media. Cluster 12 most strongly expressed the cell maintenance and efflux pump signatures, but with moderate fluctuation in the remaining pathways. The region defined by clusters 2, 4 and 10 are largely characterized by an absence, or possibly repression, of expression across most molecular processes, suggesting these cells are healthy.

**Figure 3.**
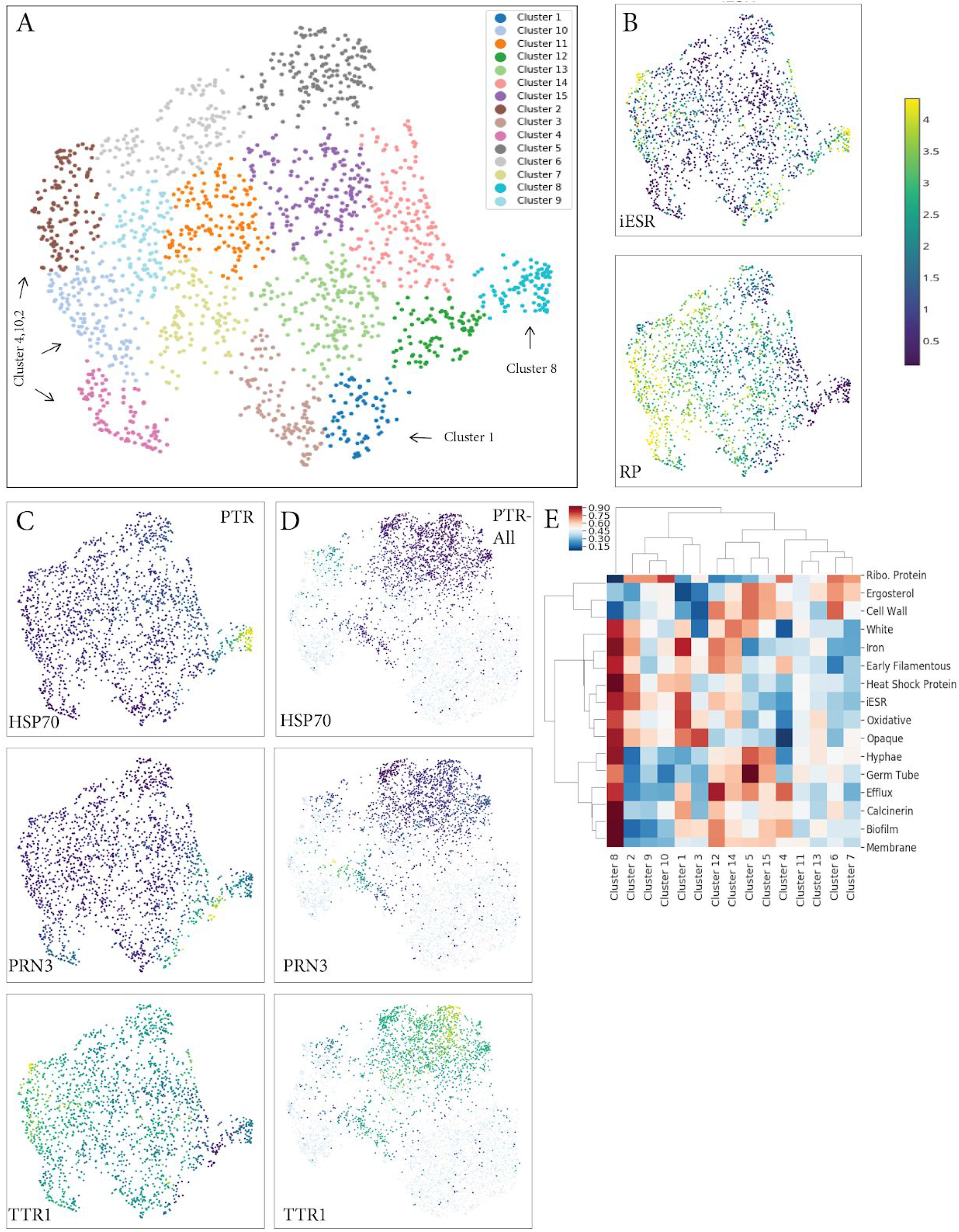
**A.** Visual representation (UMAP) of the relationships of only prototrophic (PTR) cells. In total, 15 subpopulations are highlighted via Louvain clustering. **B.** Pattern of expression of the iESR and Ribosomal Protein (RP) signatures mapped onto the UMAP embedding of A. **C.** Gene expression of the endoplasmic reticulum chaperone HSP70 (top), the ABC transporter PRN3 (middle) and dithiol glutaredoxin TTR1 (bottom) across PTR cells. **D.** Gene expression of the endoplasmic reticulum chaperone HSP70 (top), the ABC transporter PRN3 (middle) and dithiol glutaredoxin TTR1 (bottom) across all cells (drugs/timepoints) but with PTR cells highlighted in the UMAP embedding of all cells. **E.** Summary of expression of all signatures across the Louvain clusters identified across only PTR cells.

At the gene level, we observe expression of the endoplasmic reticulum chaperone HSP70 localized to cluster 8, the ABC transporter PRN3 localized to clusters 1, 12, and TTR1, a dithiol glutaredoxin, is most highly expressed in clusters 2, 4 and 10 (**Figure 3C**). We located these PTR cells which express markers HSP70, PRN3 or TTR1 back in the UMAP embedding for all cell populations (**Figure 3D**). HSP70 is most highly expressed in mustard cluster 6 of **Figure 2B**. As discussed above, this is a highly diverse clustering containing PTR cells and cells from each drug exposure. We asked what genes were strongly differentially expressed between these this cluster and remaining cells (**Supplemental Table 3)** and found that many have well established roles in different stress responses including HSP21, HGT6 and CAS5 (core stress response), GAC1, XYL2 and ADH2 in acid stress, SOD3, YCF1 and OXR1 in oxidative stress, consistent with the high expression of the iESR signature in this cluster (**Figure 2E**). Other genes are known hyphae morphology related genes including YHB1, UCF1, XYL2, FAB1 and REG1, or genes with established roles in virulence including HSP21 and YHB1.

PRN3 is mostly highly expressed in dark blue cluster 1 of **Figure 2B**, a cluster enriched for late NYS-72 survivors We asked what genes were strongly differentially expressed between these this dark blue cluster 1 of **Figure 2B** and neighbouring PTR cells of cluster 11, and identified several genes with established roles in drug resistance (eg RPL24, FMP45, ERG25), biofilm formation (eg TY37), and cell wall maintenance (eg PGA31) (**Supplemental Table 4**). High iESR expression was also detected in this cluster. TTR1 is most highly expressed in the mustard cluster 6 (where HSP70 is expressed) but also expressed in the clusters predominated by PTR cells (2,4,5,11,14) of **Figure 2B**. There was in general little evidence of expression of the iESR in these clusters. Genetically modified *C. albicans* with fluorescent markers for HSP70 (GFP) and TTR1 (RFP) appeared as distinct subpopulations (**Supplemental Figure 5**).

Together the data supports the existence of three distinct subpopulations of PTR cells, and provides preliminary evidence that these subpopulations expressed different transcriptional programs. This includes, but is not limited to, variability in iESR expression levels. The fact that high iESR expressing PTR cells co-cluster with late FCZ-72 and NYS-72 survivors is consistent with the concept that a cell that “bet hedges” is more likely to survive.

### Two distinct subpopulations are observed during the tolerance phase after treatment with fluconazole

**Figure 2A** suggests that isogenic (or near isogenic) individuals respond differentially to the same challenge. In particular, there are two distinct subpopulations of FCZ-48 survivors: clusters 9, 16, 18, 20 versus the second subpopulation in mustard cluster 6 (**Figure 2B**). To better characterize the molecular differences between the two subpopulations, we restricted attention to only FCZ-48 cells and applied Louvain clustering (**Figure 4A)**. We refer to these two clusters as Response A, and the remaining clusters as response B. Differences in iESR expression broadly characterize the two responses (**Figure 4B)**. The FCZ-48 cells in the brown and pink clusters 6 and 7 map exclusively to the highly diverse mustard cluster 6 of **Figure 2B** discussed previously.

**Figure 4.**
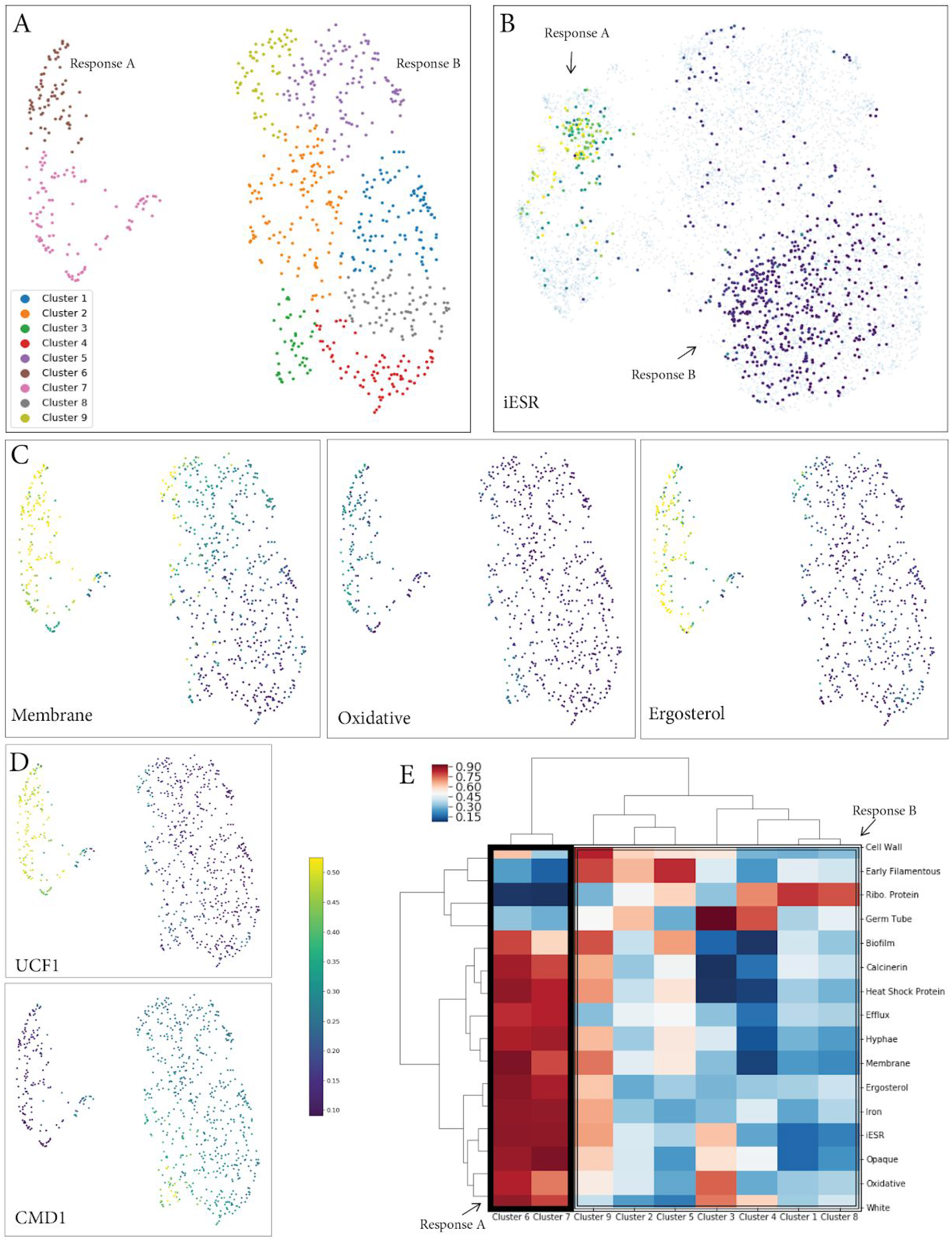
**A.** Visual representation (UMAP) of the relationships of only FCZ-48 cells. In total, 9 subpopulations are highlighted via Louvain clustering. **B.** Expression of the iESR across all cells (drugs/timepoints) but with FCZ-48 cells highlighted. **C.** Expression of the Membrane, Oxidative and Ergosterol signatures signatures mapped onto the UMAP embedding of A. **D.** Gene expression of the UCF1 (top) and CMD1 (bottom) across FCZ-48 cells **E.** Summary of all signatures across the Louvain clusters identified across all FCZ-48 cells.

Response A shows higher expression of most pathways (**Figure 4E**) including mechanisms established in the literature^9–12^ as important to fluconazole resistance including cell membrane, oxidative stress and ergosterol pathway members (**Figure 4C**).

We hypothesized that cells exhibiting response A are more likely to survive to 72 hours. To test this hypothesis, we selected markers UCF1 and CMD1 that are strongly differentially expressed between response A and B (**Figure 4D**). UCF1 (Up-regulated by CAMP in Filamentous growth 1) exhibits higher expression in Response A. Consistent with our hypothesis, down-regulation of UCF1 is associated with resistance to FCZ. Ca2+ binding protein CMD1 (Calmodulin) regulates many Ca2+ independent processes related to cellular morphology, growth and mitosis. It is not expressed in Response A and shows variable expression across Response B. We genetically modified *C. albicans* to contain fluorescent proteins (GFP for UCF1 and RFP for CMD1), grew colonies exposed to the same FCZ concentrations as was used for single cell sequencing (**Methods 3**, **Supplemental Figure 6**).

We note that CSP-48 survivors intermix with the FCZ-48 survivors in both response A and B, suggesting that although caspofungin and fluconazole are different classes of anti-fungals with distinct modes of action, patterns of heterogeneity are conserved. There is little evidence of multiple subpopulations across NYS-48 survivors with the vast majority of cells restricted to response A. Since our cell membrane signature is much higher expressed in Response A compared to B, this leads to a conjecture that cells that lowly express this process are highly sensitive to nystatin treatment as it specifically disrupts membrane function.

### Two distinct subpopulations are observed in late 72 hours survivors after treatment with fluconazole

**Figure 2A** suggests that late 72 hour survivors of fluconazole treatment partition into two distinct subpopulations. The first population resides at the convergence of clusters 3, 11 and 14, and overlap with outlier PTR cells. The second population is contained exclusively within cluster 7, and resides in close proximity to Reponse A observed at the 48 hour time point. UMAP-based visualizations of FCZ-72 identify two main subpopulations (clusters 2 and 7 versus the remaining eight clusters; **Figure 5A**). Cells from clusters 2 and 7 correspond exclusively to Response A and the remainder to Response B (**Figure 5C**). The pathways and processes differentially expressed between these two subpopulations share many similarities to the observations made in the context of 48 hour post-FCZ treatment (**Figure 5B**).

**Figure 5.**
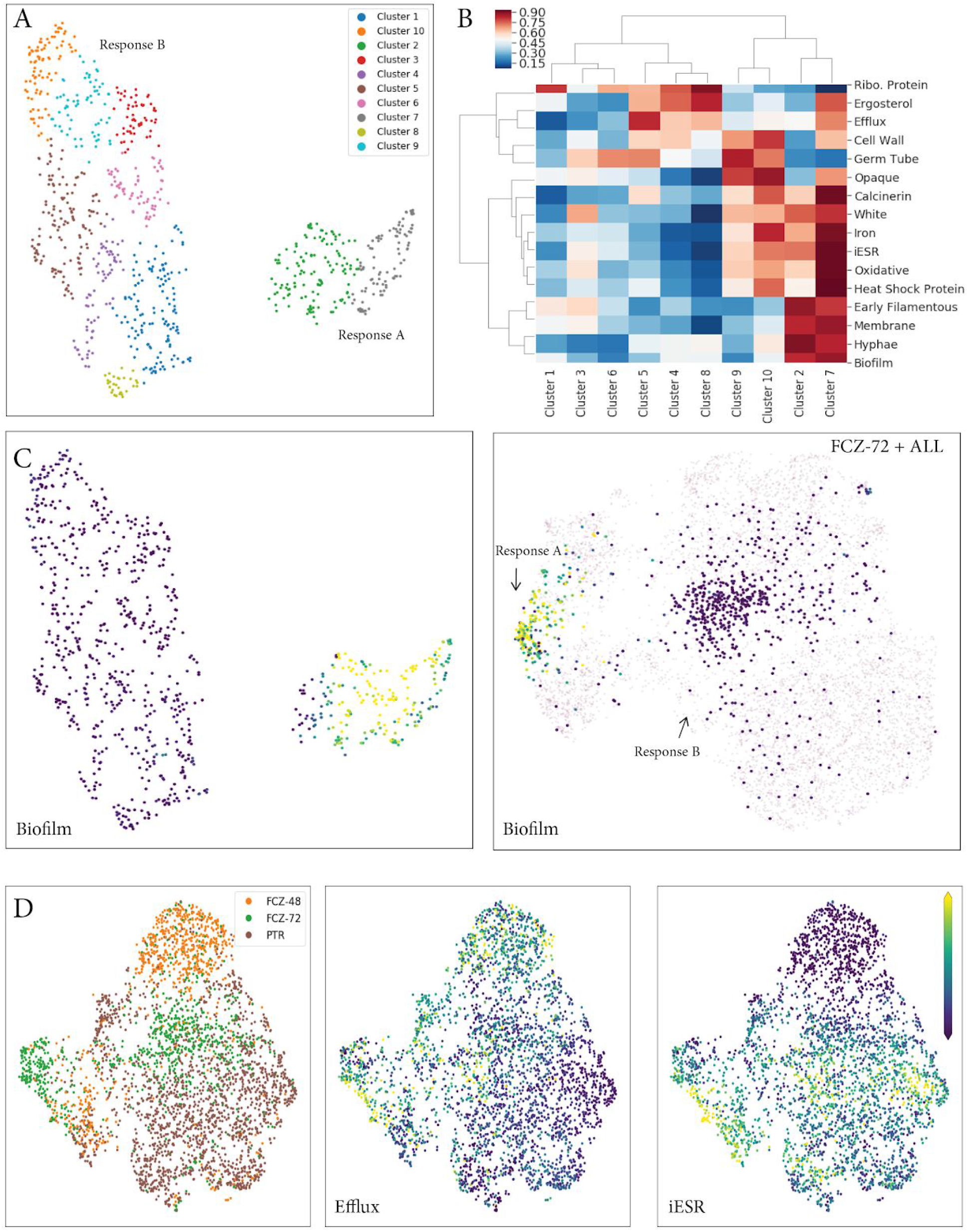
**A.** Visual representation (UMAP) of the relationships of just FCZ-72 cells. In total, 10 subpopulations are highlighted via Louvain clustering. **B.** Summary of all signatures across the Louvain clusters identified across all FCZ-72 cells. **C.** Expression of the Biofilm signature onto the UMAP embedding of A (left) and onto the UMAP embedding of all cells (drugs/timepoints) but with FCZ-72 cells highlighted (right). **D.** UMAP based visualization of the relationship between PTR, FCZ-48 and FCZ-72 cells labelled by cell of origin (left), expression of the Efflux (middle) and iESR (right) signatures.

We observe however that the FCZ-72 population closest to Response B has shifted slightly towards the expression profiles of PTR in comparison to the FCZ-48 Response B cells (**Figure 2A**). We searched our signatures for those whose expression at 72 hours more closely resembled PTR cells, in comparison to early 48 hour treatment. Both the ergosterol and efflux pathways were more lowly expressed in both FCZ-72 and PTR cells than in FCZ-48. Conversely, stress pathways including oxidative stress, heatshock and the iESR are more highly expressed in the FCZ-72 and PTR cells than in FCZ-48 (**Figure 5D**). In general, the expression changes associated with Response B are difficult to interpret; however they do suggest the tolerance phase at 48 hours may be stochastically probing different combinations of responses to survive, and survival of these cells, even if it is less likely than Response A, may involve the ablation of unnecessary cytoprotective pathways.

To examine the dynamics of these subpopulations from prototrophic, through the tolerance phase to late survivors at 72 hours, we selected two markers WH11 and YHB1 that showed differential expression between the two response and variability across the time points (**Figure 6A,B**). WH11 is expressed specifically in white-phase yeast-form cells and is similar in structure to *S. cerevisiae* GLP1, a gene coding for a plasma membrane protein involved in membrane organization and involved in maintaining organization during stress conditions. WH11 is strongly expressed in almost all PTR cells (both in Response A and B), loses expression in FCZ-48 but regains expression in FCZ-72 in Response B. The nitric oxide dioxygenase YHB1 is only expressed in cells occurring in Response A at all time points PTR, FCZ-48 and FCZ-72. Colonies exposed to the same FCZ concentrations as was used for single cell sequencing but genetically modified to express fluorescent markers for WH11 (GFP) and YHB1 (RFP) (**Methods 3**) verified that these two populations existed at 72 hours (**Supplemental Figure 7**).

**Figure 6.**
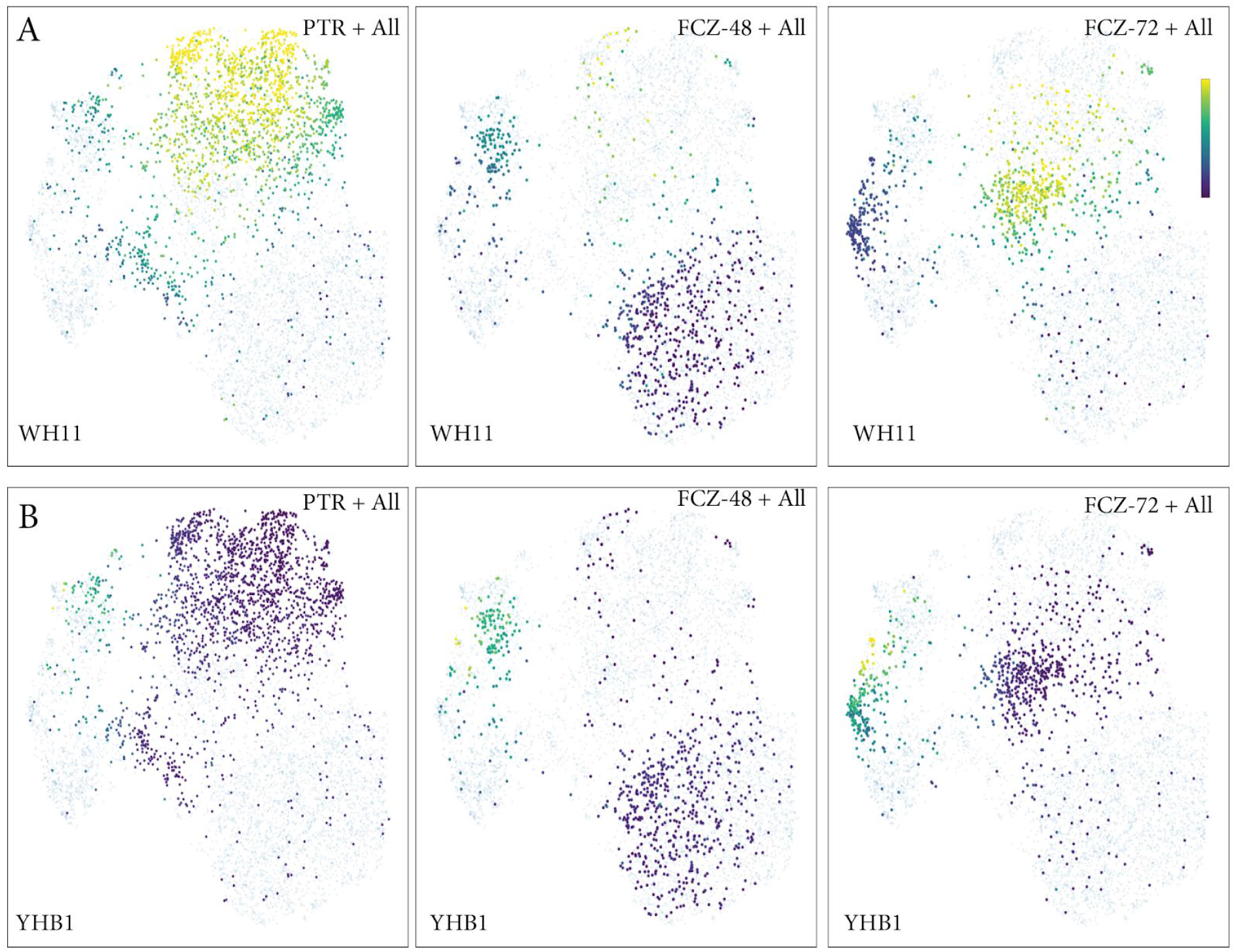
**A.** Gene expression of WH11 across all cells (drugs/timepoints) but with PTR cells highlighted (left), FCZ-48 cells (middle), and FCZ-72-cells (left) highlighted. **B.** Gene expression of YHB1 across all cells (drugs/timepoints) but with PTR cells highlighted (left), FCZ-48 cells (middle), and FCZ-72-cells (left) highlighted.

### Two distinct subpopulations are observed in late 72 hours survivors after treatment with nystatin

Late surviving NYS-72 cells exhibited evidence of transcriptional heterogeneity with two subpopulations consisting of exclusively cluster 6 or clusters 1, 17, and 19 in **Figure 2B**. Both subpopulations are contained in our so-called Response A. Due to technical reasons the NYS-48 profiles produced fewer than expected cells, although these same two Response A subpopulations are also observed during the tolerance phase.

**Figure 7A** depicts the relationships between just NYS-72 cells. Here green toned clusters 8 and 9 correspond to cluster 6 in **Figure 2B**, a highly diverse cluster that contains cells from a populations. Although there is no difference in expression of the iESR or ribosomal protein (RP) signatures, expression of the white, heatshock proteins (HSP), oxidative, iron stress and membrane signatures are all localized to cluster 6 (**Figure 7B-E**). Since UCF1 is expressed in cluster 6, we used our GFP-tagged strain of C. albicans to observe cells grown in the presence of NYS (**Supplemental Figure 8**).

**Figure 7.**
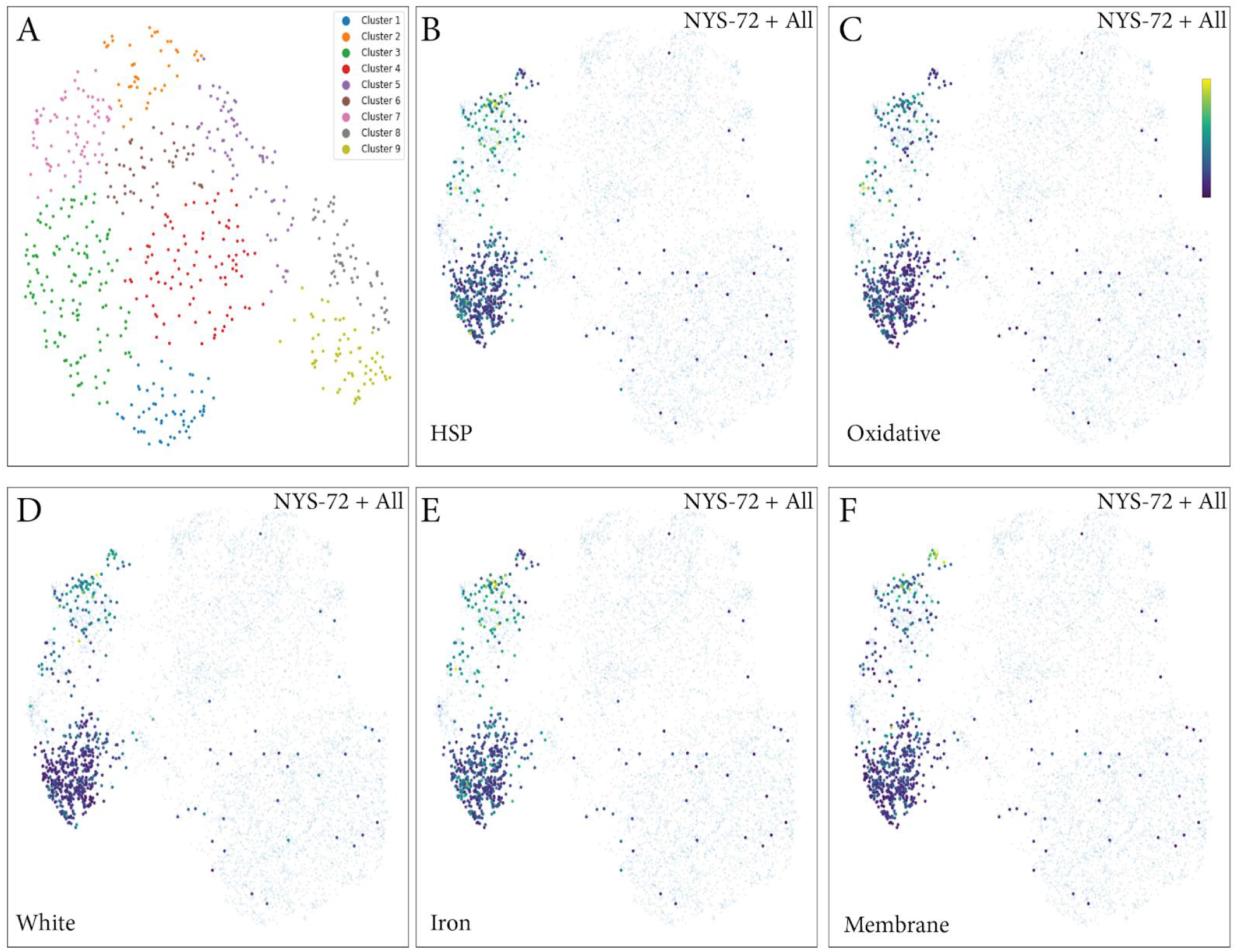
**A.** Visual representation (UMAP) of the relationships of just NYS-72 cells. In total, 9 subpopulations are highlighted via Louvain clustering. **B-F.** Expression of the Heat Shock Protein, Oxidative, White, Iron and Membrane signatures mapped onto the UMAP embedding of all cells (drugs/timepoints) but with NYS-72 cells highlighted.

## Discussion

*C. albicans* SC5314 populations, either prototrophic or grown in the presence of one of three anti-fungals, were transcriptionally profiled using a nano-litre droplet-based single cell sequencing approach optimized for the fungal setting. The prototrophic population is likely isogenic or near isogenic, thereby minimizing the chance for preexisting genetic polymorphisms that confer resistance. In terms of **Figure 1A**, the blue population rooted by * before t_0_ does not exist. In support of this, we observe heterogeneity in gene expression across the prototrophic population but do not observe distinct subpopulations with clear boundaries (**Figure 2A**), with perhaps the exception of a few cells that belong to the mustard colored cluster 6 of **Figure 2B**. Some but certainly not all of this variability is explained by variation in cell cycle. In fact, over the 15 clusters found computationally across this population, many different processes show differential expression including the induced environmental stress response (iESR), ribosomal proteins (RP) and other processes linked to drug response/stress including efflux pumps.

Microscopy confirms that the vast majority (>99%) of the cells in the prototrophic populations are of the white yeast morphology, however gene expression signatures for early germ bud and late hyphae stages do show variability. We conjecture that is is at least in part due to the manner in which these signatures were originally formed. That is, methodology from previous studies that used supervised analysis of gene expression profiles between white yeast and other morphologies in bulk populations may not be sufficiently robust to capture a presumably multi-step trajectory between the target morphologies. We conjecture that even healthy cells cycle through regulatory programs that represent proto-opaque, budding or filamentous morphologies. In other words, healthy cells “bet hedge” with various stress responses including towards a change in morphology. The most extreme example of this are those prototrophic cells that cluster with the mustard cluster 6 of **Figure 2B**. These are extremely active cells expressing many pathways simultaneously. We conjecture that prototrophic cells in the core or upper region of Figure 2A do not successfully transition to the tolerance phase (**Figure 1A**, green region B).

Each drug challenge population was profiled after 48 hours, a latency chosen to provide sufficient time for the drug to influence the colony but too short to allow genetic adaption and resistance. Survivors profiled at 48hrs under both fluconazole and caspofungin largely partition into two subpopulations, which we have termed response A and B throughout this manuscript. In broad terms, cells exhibiting response A appear transcriptionally active, expressing many stress and maintenance pathways including the proto-filamentous signatures described above. This latter statement is supported by microscopy of the populations that establish a higher fraction of filamentous cells in response A compared to response B. Perhaps response A represents an aggressive *tour de force* while response B represents a more passive cell arrest strategy.

In fact, the fluconazole and caspofungin populations intermix in both response A and B, even though they represent two distinct classes of antifungals with different modes of action (disrupting ergosterol versus cell wall biosynthesis), suggesting that cells choose a survival path that is in some cases independent of the specific drug. We are currently investigating whether this bimodal response is maintained for other-azoles and other-fungins and including additional drugs representative of all five classes of anti-fungals.

Our data suggests that 72 hour survivors, which have presumably had sufficient time to develop genetic resistance, originate from both A and B responses at 48 hours. This is apparent from both the single cell transcriptomics-based clustering and from microscopy. Given that response A individuals are more active transcriptionally, we would conjecture that this response is enriched for individuals with acquired genetic resistance (yellow type of **Figure 1A**) with response B relatively enriched perhaps for individuals who solely exploited epigenetic regulatory programs to survive the chemical insult (orange type of **Figure 1A**). We are also currently conducting experiments with pulse drug delivery to better ablate cell escape simply due to better guarantee the population is continually challenged to evolve resistance following Cowen et al.^67^ We are also exploring a greater range of concentrations including the MIC_25_, and MPC (Mutant Prevention Concentration)^68^ following EUCAST guidelines (www.eucast.org). This data, and additional whole genome sequencing (single cell or bulk) of survivor populations, would allow us to distinguish between epigenetic versus genetic survival mechanisms. The next phase of our work will be to combine the transcriptomic data with whole genome DNA sequencing to identify plausible molecular mechanisms for rapid evolution perhaps based on genomic neoplasticity and instability^18–20,22,24,25,69^.

Unfortunately technical problems in the nystatin profiles at 48 hours greatly reduced the number of successfully profiled cells, however from the paucity of nystatin 48 and 72 hour cells, there is evidence of distinct subpopulations that omits response B. We are currently adding more runs to better guarantee statistical power^70^ with our fungal single cell sequencing device for nystatin and the other drugs (including caspofungin at 72 hours). This additional data will allow us to move towards fuller pseudotime trajectories^66,71–77^, a common powerful approach in single cell studies, that frame-by-frame track changes from prototrophic through tolerant to resistant cells. The number of cells per sample and the number of distinct transcripts harvested per cell are two factors that have been challenges for pseudotime reconstructions. However, conceptually a more difficult challenge lies in the fact that expression profiles of 72 hour survivors (perhaps not surprisingly) map closer to prototrophic cells than 48 hour tolerance cells. This is in part, but not exclusively, due to the fact that the induced environmental stress response involves several hundred genes, and it is most strongly expressed in early stages.

Combination therapies use more than one drug simultaneously to reduce the probability of acquiring resistance, permit the use of lower levels of each individual drug, and improve treatment specificity^78–80^. They are now used in many contexts, however, identifying synergizing compounds is challenging, and mechanistic explanations are difficult to establish^81^. A molecular level understanding of drug tolerance in *C. albicans* will open a door to a long corridor that ends in a new type of therapeutic that targets the tolerance phase. By careful examination of the trajectories across different conditions, drugs and concentrations, we can identify the most likely “cut-points” along these paths, that when targeted, would eliminate the grace period for an individual to acquire resistance.

## Acknowledgements

We thank L Pachter and the organizers of BIRS workshop #17w5134 (Oaxaca, Mexico) for several conversations that motivated this effort especially w.r.t. affordable open biotechnology. We thank CA Jackson and D Gresham for early access to their protocol, C Law for assistance with the microscopy, A Villani for several nice optimizations, and members of the McCarroll lab for their careful advice and outstanding responsiveness.

## Data Availability and Reproducibility

Jupyter/R code and single cell transcriptome data is available from the authors’ upon request within a collaborative context while this work remains a preliminary manuscript.

## Methods

### 1. Nanolitre droplet-based single cell RNA-sequencing (DROP-seq) for Fungi

Yeast (*S. cerevisiae*) has previously been examined by single cell sequencing using the Fluidigm C1 system but this handles less than 100 cells^41^. Jackson et al.^42^ sequenced ~40K *S. cerevisiae* cells with the commercial Chromium (10X Inc.) system. We opted to build a fungal DROP-seq modified from the original approach presented in Macosko et al^43,44^, to address issues of cost and flexibility in comparison with commercial alternatives. In general, DROP-seq devices have been shown to be near equivalent to commercial systems^45^. In particular, here we built a printed circuit board to control microfluidic flow inspired by Stephenson et al^46^, and 3D printed plastic syringe pumps and cheap cameras inspired by Booeshaghi et al^47^ (**Figure 1A**). The cost per device is well below $1K USD.

### 2. Strains and media

*C. albicans* SC5314 cells were grown in YPD liquid media (2% D-glucose, 2% peptone, 1% yeast extract, 0.01% uridine) and incubated at 30°C for 12-16 hours. Afterwards an aliquot of 10^8^ cells was taken and used as the prototrotrophic sample. Then, we introduced 1ml RNAlater (Sigma # R0901) and froze the resultant colony at −20°C for later use in the DROP-seq. Other aliquots were used for drugs treatment experiments. In order to have enough log-phase cells for multiple drugs at different timepoints, we performed the following. Cells were pelleted and resuspended in 1ml of YPD. Then, 250μl of this suspension was combined with 15ml of fresh YPD and placed in a shaker incubator at 30 °C for 4-5 hours. Finally, on the order of 10^8^ of these cells were placed in 10ml of YPD. Each suspension was then subjected to drug treatment.

### 3. Anti-fungal drug treatment

A concentration of 0.01 mg/ml was chosen for both fluconazole (Sigma #F8929) and nystatin (Sigma #N6261), representing a moderate dosage relative to their reported MIC50 levels^8,10,48–50^. A concentration of 1ng/ml was used for caspofungin (Sigma #SML0425), a compound that interrupts cell way biosynthesis^51–53^; this is well below its reported MIC50 levels and chosen in order to ensure a sufficient number of survivors to generate single cell profiles. The target drug was delivered to the individual colonies from step 2 and incubated at 30 °C for 48 or 72 hours. Cells at these time points were strained (pluriStrainer® 20 μm) and collected in fresh tubes. This was done in order to minimize the likelihood that the microfluidic chip would block due to large hyphae and pseudohyphae morphologies. We observed that germ tubes up to four times the length of the mother cell can still be processed for drop-seq analysis (**Supplemental Figures 3, 4**). Such cells are well within the hyphal transcriptional profile^54^. This suggests that our results do contain profiles of hyphae and pseudohyphae cells.

### 4. Spheroplasts

The *C. albicans* setting required an optimized protocol for cell preparation with specific techniques to remove the cell wall and induce stable spheroplasts. Towards this end, we experimented with different concentrations of zymolyase (0.1, 0.2 and 0.4U zymolyase (BioShop # ZYM002), to 10^7^ cells in 100 ul of sorbitol 1M) at different time points (stored at 37 °C for 10, 20, 30 mins) before processing with the DROP-seq. To compare against untreated populations via microscopy (Leica DM6000), cells were stained with calcofluor white. We concluded that concentrations in the range 0.1-0.2U after 20 minutes are able to induce spheroplasts that remain sufficiently stable for processing with our DROP-seq.

### 5. Cell preparation

Drug treated colonies at either the 48 or 72 hours were pelleted by brief centrifugation and washed two times with 1ml RNAlater. Cells were then resuspended in 0.5ml of RNAlater and stored at room temperature for one hour. Afterwards, they were put in −20 °C for at least 24 hours before passing through the DROP-seq.

At the time of the DROP-seq, an aliquot of 10^7^(OD=0.68 in 660 nm) cells from each colony, and wash each colony three times with sorbitol 1M. The cells are then resuspended in 100μl sorbitol 1M + 0.2 U Zymolyase and incubated at 37 °C for 20 minutes (as per our findings in Methods 3). Next, the cells were pelleted and resuspended again in 0.5ml of cold and fresh RNAlater for five minutes. Now, the cells were washed (centrifuged and pelleted) with 1ml of washing buffer (1M sorbitol, 10mM TRIS pH 8, 100ug/ml BSA) three times. Finally, 10^6^ cells (OD=0.08 in 660nm) were resuspended in 1.2 ml of washing buffer. This cell suspension was then used as input to the DROP-seq device.

### 6. Fungal DROP-seq protocol

Cell preparation generally follows the protocol given by Macosko et al.^43,44^ with some exceptions. Whereas Macosko et al recommends a ratio of 100K cells to 120K beads for DROP-seq, we found that a ratio of 1M cells for 120K beads generated a sufficient yield of cDNA as per the Agilent Tapestation. Jackson et al.^42^ used 5M cells as input to the Chromium (10X Inc.) system. Furthermore, whereas Macosko et al. use 1ml of lysis buffer, we used 1.2 ml, and instead of 13 PCR cycles, we used 17 (Jackson et al. used 10 cycles). Samples were sequenced using the Illumina NEXT-seq following a standard protocol^43,55^ (200M read/sample).

### 7. Bioinformatics and statistics for the single cell profiles

In general, all computations were performed using Python version 3.67 or R version 3.6.1. Gene abundances were estimated from raw sequencing data using the end-to-end pipeline Alevin^56^ which optimizes UMI deduplication and reduces the number of discarded (multimapped) reads. SCANPY^57^, a python-based toolkit for analyzing single-cell gene expression data was used for data quality control and preprocessing. We selected cells with at least 30 genes and 50 read counts under the condition that less than half were found in ribosomal genes (RDN). We removed genes that expressed in less than 20 cells. Normalization, imputation and batch correction were performed by scVI^58,59^, a tool which implements a probabilistic model of mRNA capture and uses a variational autoencoder to estimate priors across batches and conditions (**Supplemental Figure 1**).

### 8. Novel approaches for identifying subpopulations with distinct states and expression patterns

To identify subpopulations of cells with similar gene expression patterns in an unsupervised manner, dimensionality reduction and visualization were based on UMAP^60^ with Louvain^61^ clustering. To assist in the identification of trajectories within and between conditions, we developed a novel graph variational autoencoder (VAE) (**Supplemental Methods 1**) to reduce dimensionality using Louvain61 clustering. Additionally, a novel measure of complexity for single-cell gene expression profiles was developed (**Supplemental Methods 2)** for the identification of cell types and states. The resultant latent space can then by readily visualized.

### 9. Gene signatures

**Supplemental Tables 2, 5** lists all of the gene signatures used throughout the analysis. In some cases, gene signatures from the literature were derived in other organisms and required orthology mappings to *C. albicans*. This includes the *S. cerevisiae* derived Environmental Stress Response (ESR)^40^ containing 859 genes and divided into three broad categories called the induced ESR (iESR; genes that are differentially regulated in response to environmental xenobiotics, conditions or other challenges), the ribosomal proteins (RP) and the ribosomal biogenesis genes (RiBi; involved in rRNA production, growth and cell division). To generate a *C. albicans* version of the ESR, we downloaded *C. albicans* (strain SC5314) assembly 21 and Sc (S288C) orthology maps from the Candida Genome Database (http://www.candidagenome.org/), and synteny maps from the Candida Gene Order Browser^62^ at this website. For each *S. cerevisiae* gene we almost always used synteny as the primary attribute determining the correct *C. albicans* orthologue. When these databases failed to identify a *C. albicans* geene, we manually evaluated the quality of the reciprocal best BLAST-protein alignment between *S. cerevisiae* and *C. albicans*. In total, orthologs for 642 of the 859 *S. cerevisiae* ESR genes were identified (**Supplemental Table 5**).

The list of gene signatures consists of the following: ras pathway^34^, ergosterol biosynthesis^12^, hyphae morphology^29^, calcineurin pathway^82^, oxidative stress^83^, cell wall biosynthesis^51,84^, efflux pumps^10,11^, pseudohyphal morphology^85^, heat shock^16^, biofilm^36^, opaque morphology^58^, iron starvation^51,40^, parasexual and meiosis^26,27^, germ tubes^86^, white morphology and early filamentous morphology^31^.

### 10. Novel approaches for exploring the molecular components of subpopulations

Our univariate analyses started with a simple Welch t-test to identify genes that are strongly differentially expressed between two given target populations. When selecting for marker genes in the downstream microscopy validation studies, we narrowed our focus towards genes that were strongly up-regulated in one cluster. Our multivariate analyses started with the VISION tool^63^ to identify sets of genes that are strongly differential between two given target populations. Given a gene signature (**Methods 9**), VISION computes a signature score based on a combination of gene expression and a precomputed cell-cell similarity map. We also developed a novel Generative Adversarial Network (GAN) (**Supplementary Methods 3**) to explore gene association networks in our data.

### 11. Live cell imaging

Live cell imaging was used to validate subpopulations identified with the single cell transcriptional profiles. We proceeded as follows.
11a. Genes were selected whose expression profiles were differential expressed between subpopulations at each time point as described in **Methods 10**. This list of genes was winnowed down to one gene in each subpopulation with high expression levels. We limited validation studies to the two most distinct subpopulations at each time point (prototrophic, FCZ 48h, FCZ 72h). In this manner, we required three pairs of marker genes, one tagged with GFP and with this FRP.
11b. Primers were designed for these six target genes (**Supplemental Table 1**). Strain SN76(*his1Δ/his1Δ, arg4Δ/arg4Δ, ura3Δ/ura3Δ*) was chosen for gene tagging, since it is a derivative strain of SC5314 but with multiple auxotrophic markers. These markers allow for convenient selection of cells successfully transformed as the target genes with integration the fluorescent protein gene in addition to the auxotroph marker (eg *HIS1*) via homologous recombination.
11c. Benchling (https://benchling.com) was used to design the sgRNAs and we followed the CRISPR/Cas9 protocol with the plasmid pV1093 from Min et al^64^. This includes two PCR reactions to fuse the SNR52 promoter to the sgRNA scaffold and terminator. The third PCR reaction amplifies the final sgRNA cassettes. Two different plasmids pENO1-iRFP-NATr (Addgene Inc) and pFA-GFP-HIS1 were used to design the repair segment. The construction of the Cas9 cassette proceeded as per Min et al. Amplification of the the Cas9 cassette with PCR used the following schedule: 98°C for 3 minutes, 98°C for 30 seconds, 63°C for 30 seconds, 72°C for 5 minutes and 30 seconds. Steps 2 to 4 have been repeated for 34 rounds followed by 72°C for 10 minutes and finally the reaction finished in 4°C. The repair DNA must be amplified with the designed primers described in **Supplemental Table 1** in 8-12 PCR tubes with 0.1μl plasmid (500ng/ml), 2.5μl forward primer, 2.5μl reverse primer, 1μl 10mM dNTP, 33.65μl nuclease free water, 10μl 5X HF PCR buffer and 0.25μl phusion polymerase in each tube.
11d. Preparation of cell colonies for microscopy. Cells that were successfully transformed were grown and harvested for each drug at each timepoint in a manner identical manner to that used for the single cell experiments (**Methods 4**). At time of microscopy, cells were collected, washed with H2O and transferred to minimum media to minimize the background noise from normal YPD media. After, cells were mounted onto the uSlide and imaged with Nikon Ti microscope.

---

**Supplemental Figure 1.**
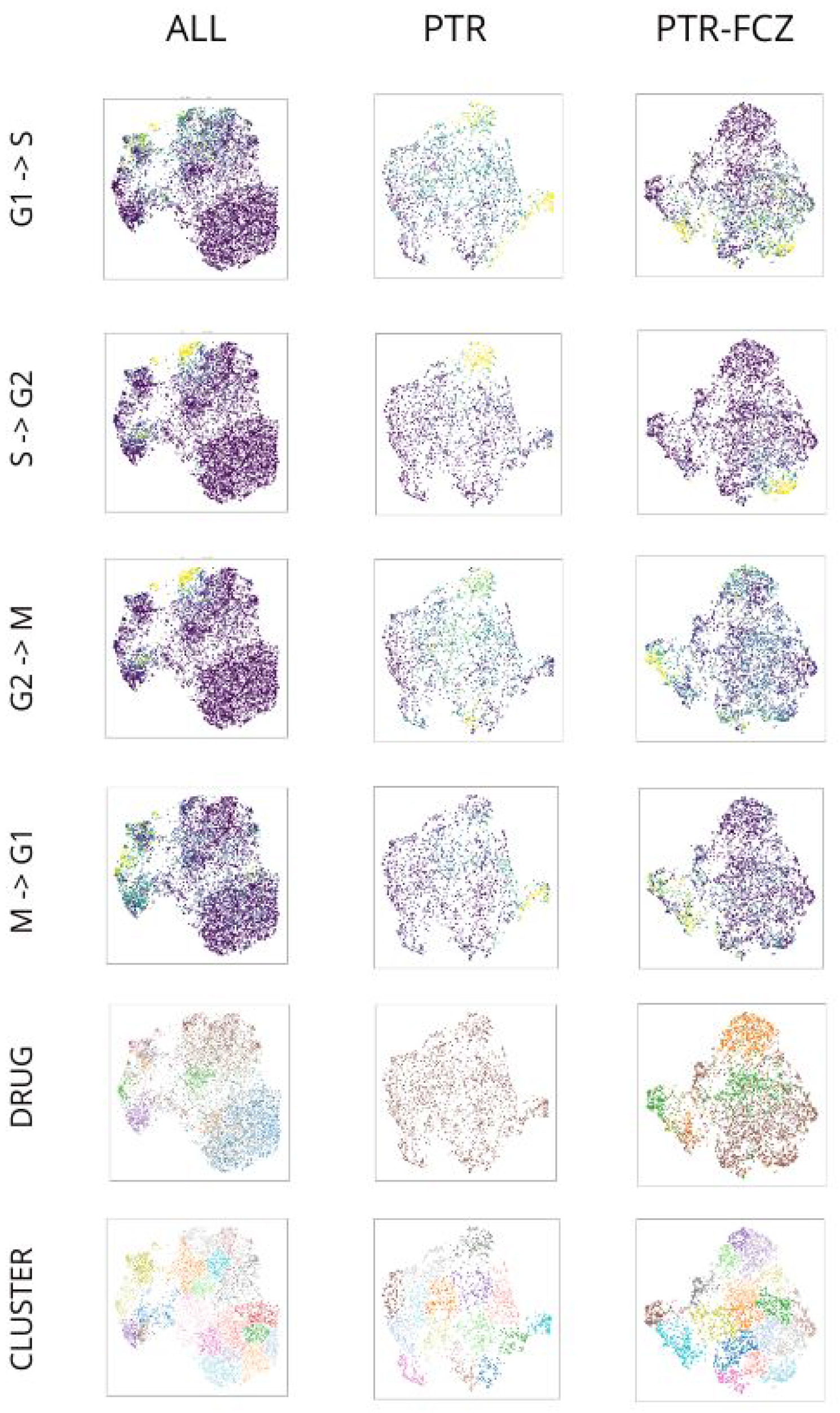
Patterns of expression of cell cycle signatures mapped onto the UMAP embedding of all cells (left), mapped onto the UMAP of PTR cells (middle), and mapped onto the UMAP of PTR, FCZ-48 and FCZ-72 cells (left).

**Supplemental Figure 2.**
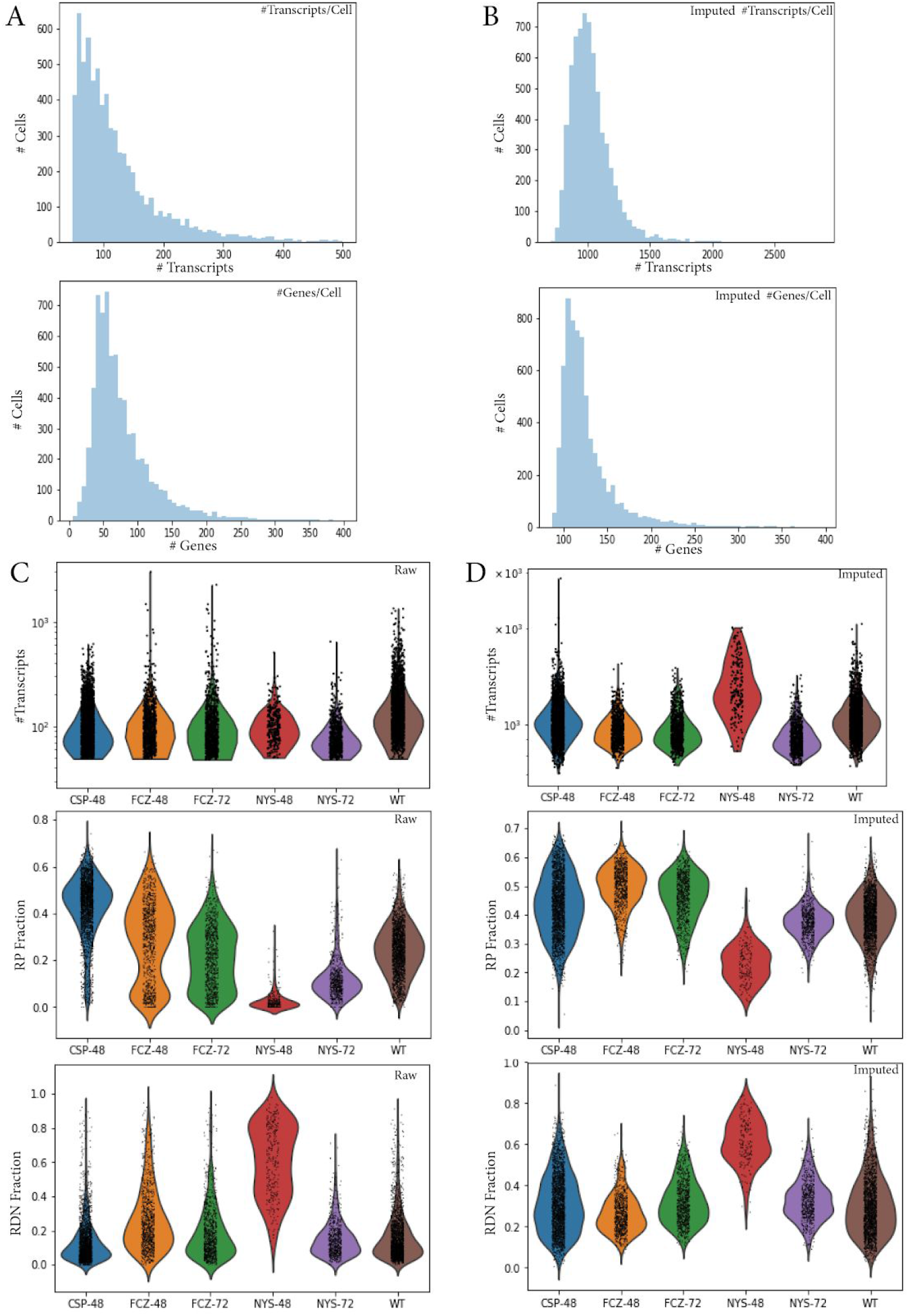
**A.** Histograms describing the number of cells with observed levels of transcripts (top) and genes (bottom) before normalization and imputation. **B.** Histograms describing the number of cells with observed levels of transcripts (top) and genes (bottom) after normalization and imputation. **C.** Violin plots describing the distribution of transcript number (top), fraction of transcripts of genes with an RP prefix (middle), and fraction of transcripts of genes with an RDN prefix (bottom) before normalization and imputation. **D.** Violin plots describing the distribution of transcript number (top), fraction of transcripts of genes with an RP prefix (middle), and fraction of transcripts of genes with an RDN prefix (bottom) after normalization and imputation.

**Supplemental Figure 3.**
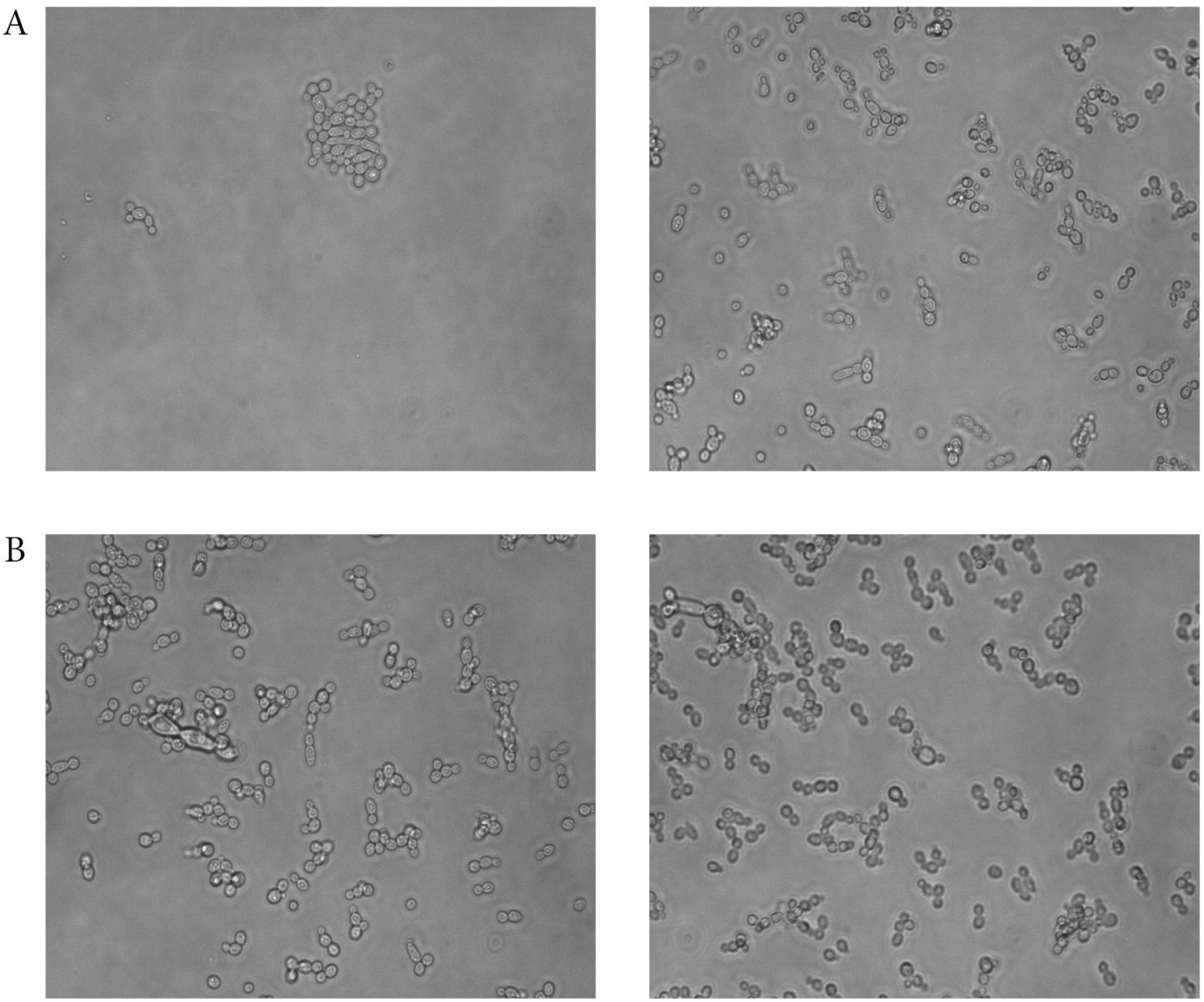
**A.** Images taken under the microscope of CSP-48 cells without (left) and with a cell strainer (right). **B.** Images taken under the microscope of FCZ-48 cells without (left) and with a cell strainer (right).

**Supplemental Figure 4.**
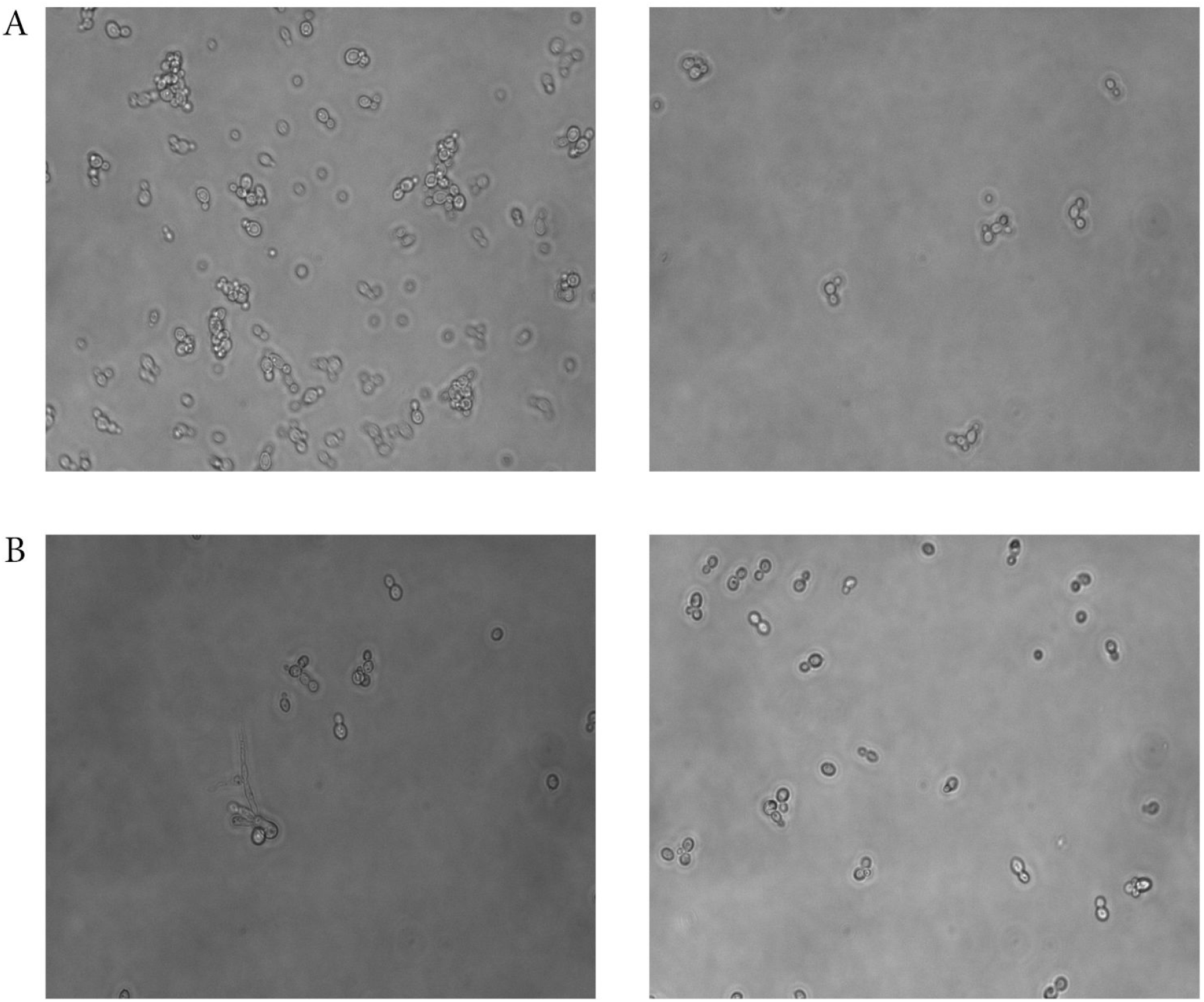
**A.** Images taken under the microscope of NYS-48 cells without (left) and with a cell strainer (right). **B.** Images taken under the microscope of FCZ-72 cells without (left) and with a cell strainer (right).

## Supplemental Tables

**Supplemental Table 1.**
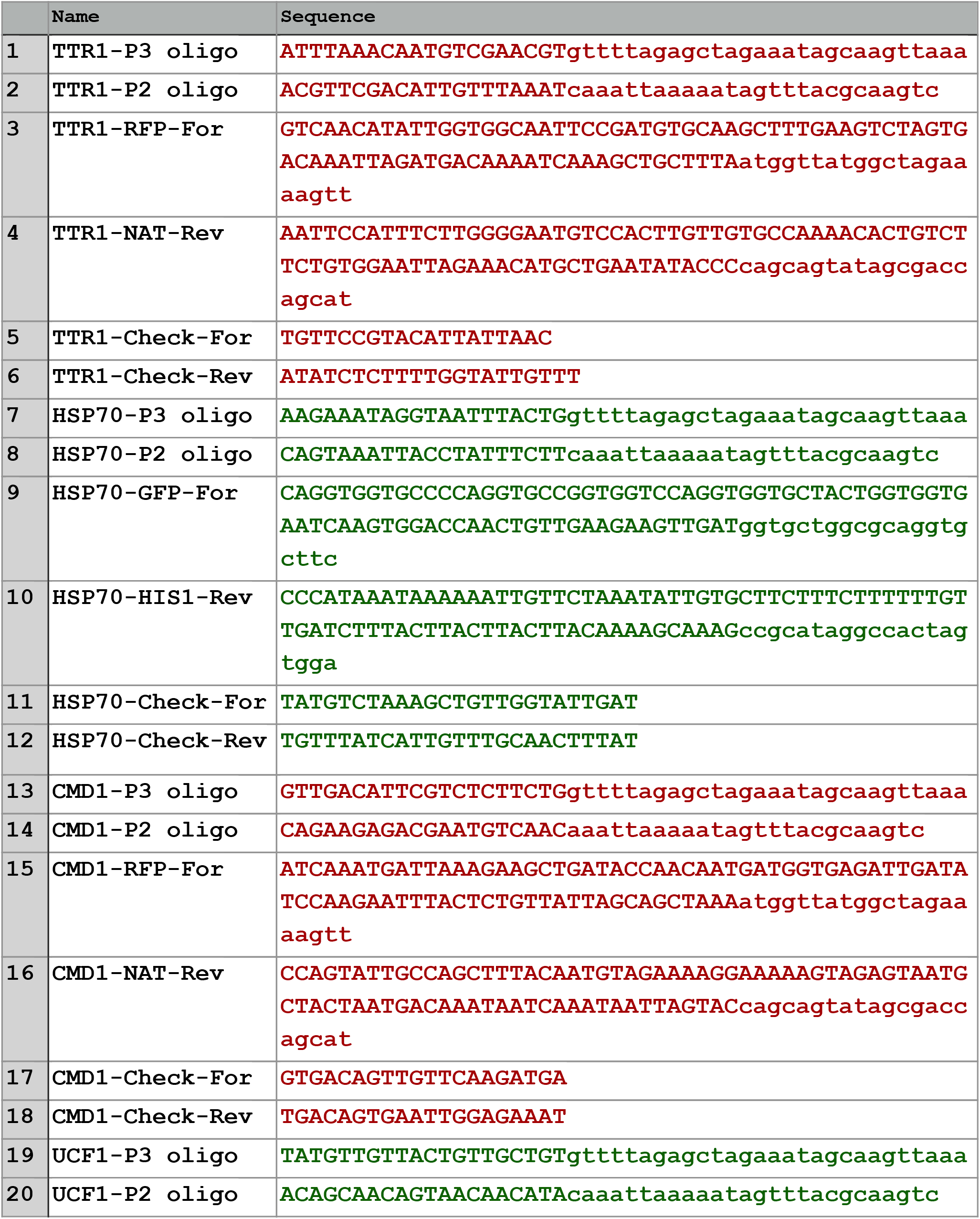
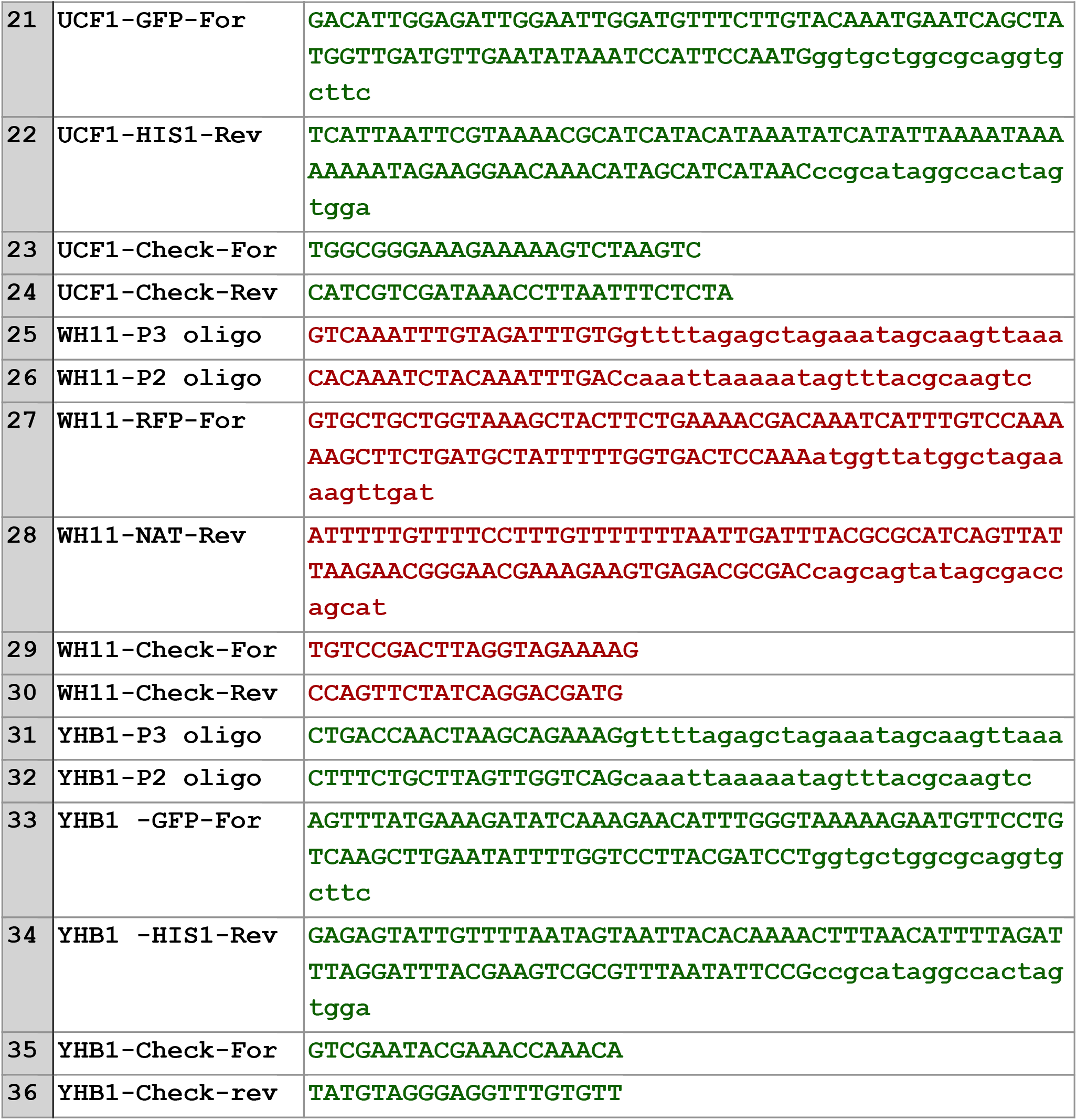
The nucleic acid sequences of forward (For) and reverse (Rev) primers used for the target genes in the live cell imaging experiments. sgRNAs (capital letters) are fused to P2 and P3 oligos.

**Supplemental Table 2.**
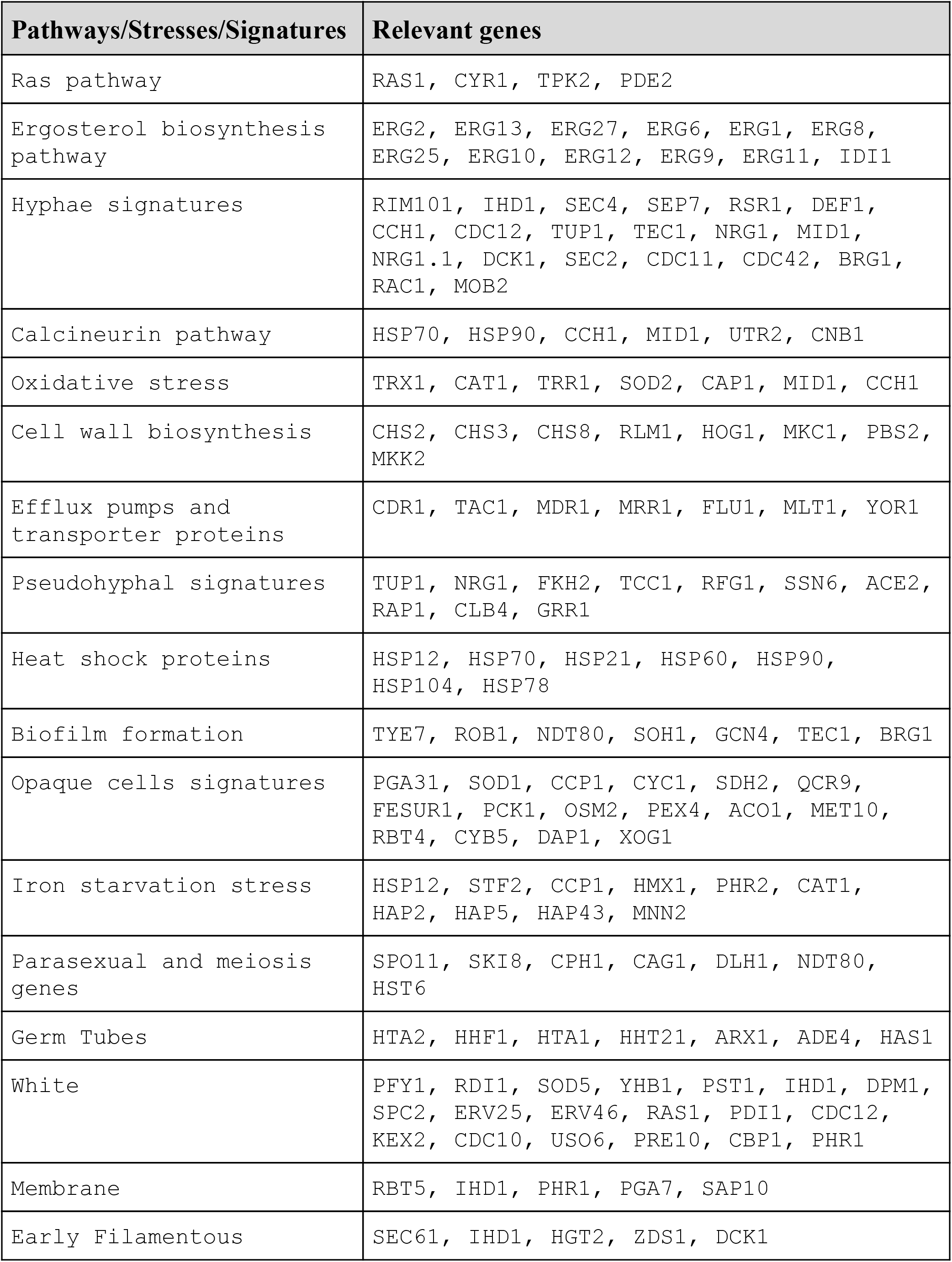
Gene sets representing pathways, processes and morphological states in *C.albicans.*

**Supplemental Table 3.** Genes which are showing strong differential expression between cluster 8 versus all the remaining clusters within the population of PTR cells.

|  | Genes |
| --- | --- |
| Log FC > 2 | HSP21, CTR1, GAC1, PGA48, FGR41, YHB1, UCF1, HSP70, TYE7, SOD3, SMC6, NAD, BLP1, HGT6, XYL2, ADH2, FAB1, PGK1, YVC1, IFE2, YCF1, OXR1, GAC1, ITS1, CAS5, APE3, REG1, NDH51, ARP9, HHT21 |
| Log FC < -2 | RPL29, RPL24A, RPL43A, RPL37B, RPL30, RPS21B, NHP2, UBI3 |

**Supplemental Table 4.**
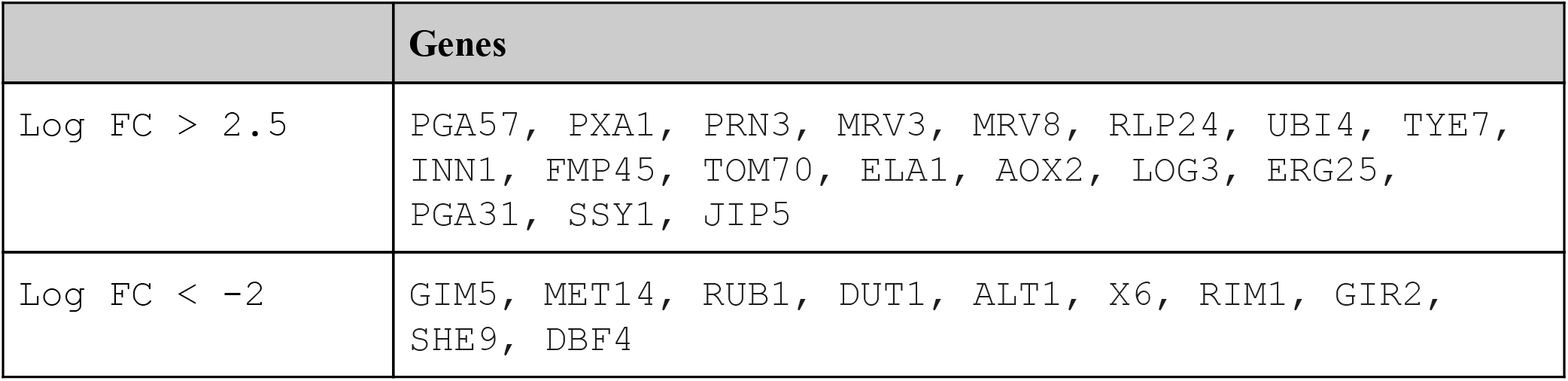
Genes which are showing strong differential expression in cluster 1 and cluster 11 of the UMAP embedding of all cells (drugs/timepoints).

|  | Genes |
| --- | --- |
| Log FC > 2.5 | PGA57, PXA1, PRN3, MRV3, MRV8, RLP24, UBI4, TYE7, INN1, FMP45, TOM70, ELA1, AOX2, LOG3, ERG25, PGA31, SSY1, JIP5 |
| Log FC < -2 | GIM5, MET14, RUB1, DUT1, ALT1, X6, RIM1, GIR2, SHE9, DBF4 |

**The excel sheet is entitled Supplemental Table 5 is available for download.**

**Supplemental Table 5.** *S. cerevisiae* ESR ortholog genes in *C.albicans.* RP(Ribosomal Protein), RiBi(Ribosome Biogenesis), iESR(involved in Environmental Stress Response)

